# Acoustically driven cortical delta oscillations underpin perceptual chunking

**DOI:** 10.1101/2020.05.16.099432

**Authors:** J.M. Rimmele, D. Poeppel, O. Ghitza

## Abstract

Oscillation-based models of speech perception postulate a cortical computational principle by which decoding is performed within a window structure derived by a segmentation process. In the syllable level segmentation is realized by a theta oscillator. We provide evidence for an analogous role of a delta oscillator at the phrasal level. We recorded MEG while participants performed a target identification task. Random-digit strings, with phrasal chunks of two digits, were presented at chunk rates inside or outside of the delta range. Strong periodicities were elicited by acoustic-driven chunk rates inside of delta, in superior and middle temporal areas and speech-motor integration areas. Periodicities were diminished or absent for chunk rates outside of the delta range, closely in line with behavioral performance. No periodicities were observed for top-down driven chunking conditions. Our findings show that phrasal chunking is correlated with acoustic-driven delta oscillations, expressing anatomically specific patterns of neuronal periodicities.

## Introduction

In written text, spaces and punctuation divide words and define phrase boundaries. In contrast, naturally spoken language is a stream of connected sounds. Embedded in the acoustic stream is information analogous to spaces and punctuation rules (e.g. intonation, stress, pauses), broadly termed ‘accentuation,’ that is used by listeners to indicate boundaries in speech associated with linguistic units. This marking is obtained by segmentation – a process by which the input signal is partitioned into temporal segments, or chunks, that are ultimately linked to a variety of linguistic levels of abstraction, ranging from phonetic segments to syllables to words and, ultimately, prosodic and intonational phrases. The segmentation process works on intervals associated with syllables (up to ∼250 ms) and on the phrasal level (0.5–2 s). For syllable-level segmentation, the neuronal mechanisms are widely studied, but they are largely unknown at the phrasal level. Only after the signal has been segmented can *effective* decoding proceed (Doelling et al., 2014; Ghitza, 2017; Ghitza and Greenberg, 2009). If a signal is incorrectly segmented, it is more difficult to form a match with internal linguistic patterns associated with syllables, words and phrases.

Perceptual segmentation, or chunking, at the syllabic level is by and large a pre-lexical process. Oscillation-based models propose that this segmentation is realized by flexible theta oscillations aligning their phase to the input syllabic rhythm (‘speech tracking’), where the theta cycles mark the speech chunks to be decoded. This is argued to function because the mean period of theta oscillations closely aligns with the mean rate of speech and the mean duration of syllables crosslinguistically (e.g. Ding et al. 2017; Varnet et al. 2017, Poeppel & Assaneo 2020). In particular, oscillation-based models of speech perception postulate a cortical computation principle by which decoding is performed within a time-varying window structure, synchronized with the input on multiple time scales (e.g., Poeppel, 2003; Ahissar and Ahissar, 2005; Lakatos et al., 2005; Ding and Simon, 2009; Ghitza and Greenberg, 2009; Giraud and Poeppel, 2012; Peelle and Davis, 2012; Gross et al., 2013). The window structure is obtained by segmentation, a process by which the input signal is partitioned into temporal chunks––windows––that are ultimately linked, by a decoding process, to different linguistic levels of abstraction. Only after a signal has been segmented can effective decoding proceed. These models proved to be capable of explaining a range of counterintuitive psychophysical data (e.g., Ghitza and Greenberg, 2009; Ghitza, 2011, 2012, 2014) that are hard to explain by conventional models of speech perception. Doelling et al. (2014) provided MEG evidence for this computation principle, showing that perceptual segmentation on the syllabic level appears to *require* acoustic-driven theta neuronal oscillations (see also: Park et al., 2015; Oganian and Chang, 2019).

At the phrase level, phrase rhythm can affect parsing (see also: Deniz and Fodor, 2019; Hilton and Goldwater, 2020; Martin, 2015). To be sure, at the level of phrases (or larger units in utterances, i.e. sentences), there is considerable variation in duration, and therefore any rhythmicity is relatively variable. There have been various studies aiming to quantify phrase length and rhythmicity, for example in English, and research by Jaffe and colleagues as well as results summarized in Auer et al. (Auer et al., 1999) suggest that typical intonational phrases are a little bit under one second in duration. The extent to which phrase level rhythmic structure supports segmentation and structural parsing is not widely studied; but insofar as one can track this level of temporal regularity, phrasal chunking can support further processing steps. Perceptual chunking at this level comprises (at least) two distinct processes: a bottom-up, acoustically-driven segmentation process (likely caused by prosodic/intonational accentuation cues) on the one hand, and a top-down, contextual parsing process (based, e.g., on syntactic cues), on the other. (We distinguish between segmentation and parsing. Segmentation results in temporal partitioning of the acoustic stream. Parsing refers to the division of the incoming signal into candidate constituents using their syntactic and semantic roles.) What kind of neuronal mechanism can realize these processes?

Delta oscillations (∼1-3 Hz) are presumed to be at the core of these mechanisms, because their frequency range corresponds roughly to the phrasal time scale (Ding et al., 2016; Keitel et al., 2018; Kösem and van Wassenhove, 2017; Mai et al., 2016; Meyer, 2017; Meyer et al., 2016). Behaviorally, it has been shown that performance in a digit retrieval task is impaired when the phrasal presentation-rate is outside of the delta range, suggesting a role of acoustic-driven delta in phrasal segmentation (Ghitza, 2017). Electrophysiologically, a recent EEG study suggested that prosody and syntax can interact in eliciting delta-band activity (Meyer et al., 2016). In another study, Ding et al. (2016) provided MEG evidence that delta activity – corresponding to the phrasal-level linguistic structure – were elicited without any acoustic cues at these time scales, interpreted there as a top-down consequence. These studies conjecture that a context-invoked delta is generated by a decoding process, guided by past and ongoing linguistic content which, in turn, plays a role in linguistically-guided parsing. Little is known about the origin of acoustic- and context-invoked delta oscillations. They may recruit distinct brain areas (Kaas and Hackett, 2000; Rauschecker and Tian, 2000), including auditory cortex, motor cortex and higher-level areas involved in linguistic processing (Ding et al., 2016; Keitel et al., 2017; Meyer, 2017; Morillon et al., 2019)(see also: Rimmele et al., 2018).

Here we focus on the cortical mechanism that may be involved in *acoustic-driven* segmentation at the phrasal level. We test the hypothesis that the decoding process on the phrasal level is guided by a flexible, acoustic-driven delta oscillator locked to phrase-size acoustic cues (Ghitza, 2017). We recorded MEG data while participants performed a digit retrieval task as in Ghitza (2017). Digit strings were used in order to minimize context-related cues. The digits in the string were grouped into pre-determined chunk patterns, with the chunk rates either inside versus outside or marginal to the delta frequency range. (See illustration of the stimulus waveforms in the Materials and Methods section.) The experiment addresses three questions: (1) Do elicited delta neuronal oscillations correlate with behavior, such that impaired performance in digit retrieval occurs if the chunk rate is outside the delta range? (2) Where in the auditory pathway do those neuronal oscillations occur? (In early auditory cortex? In ventral or dorsal auditory streams?) (3) Are these neuronal oscillations acoustically driven or context invoked?

The data show that in superior and middle temporal areas, robust neural periodicities were elicited by acoustic-driven chunk rates inside of the delta range - but were diminished when the chunk rate was outside of the delta range, in line with behavioral performance. In speech-motor integration areas, periodicity was present albeit clearly diminished even for chunk rates inside the delta range. No periodicities were evident for top-down chunking (no-chunk) conditions. In contrast, strong theta responses were elicited for all conditions, including the no-chunk control condition, reflecting the single digit presentation rate. The observed delta band periodicities, in a hierarchy of brain areas, were acoustically driven and not context driven.

## Results

### Behavioral results

Dprime scores were the highest in the 1.8Hz condition (mean = 2.19, SD = 0.50) (Fig. 1), i.e., when the chunk rate is inside the delta frequency range. Lower dprime scores were registered in the 2.6Hz condition (mean = 1.74, SD = 0.43), when the chunk rate was just outside the delta range. The lowest scores were registered in the no-chunk condition (mean = 1.38, SD = 0.30). The difference in scores was significant between all combinations of condition pairs (1.8Hz condition vs. 2.6 Hz: p = .0079 ; 1.8 Hz vs. no-chunk: p = .0008; 2.6Hz vs. no-chunk: p =.0048; Bonferroni p-corr = .0167), with highest performance for chunk rates inside the delta range.

**Fig. 1.**
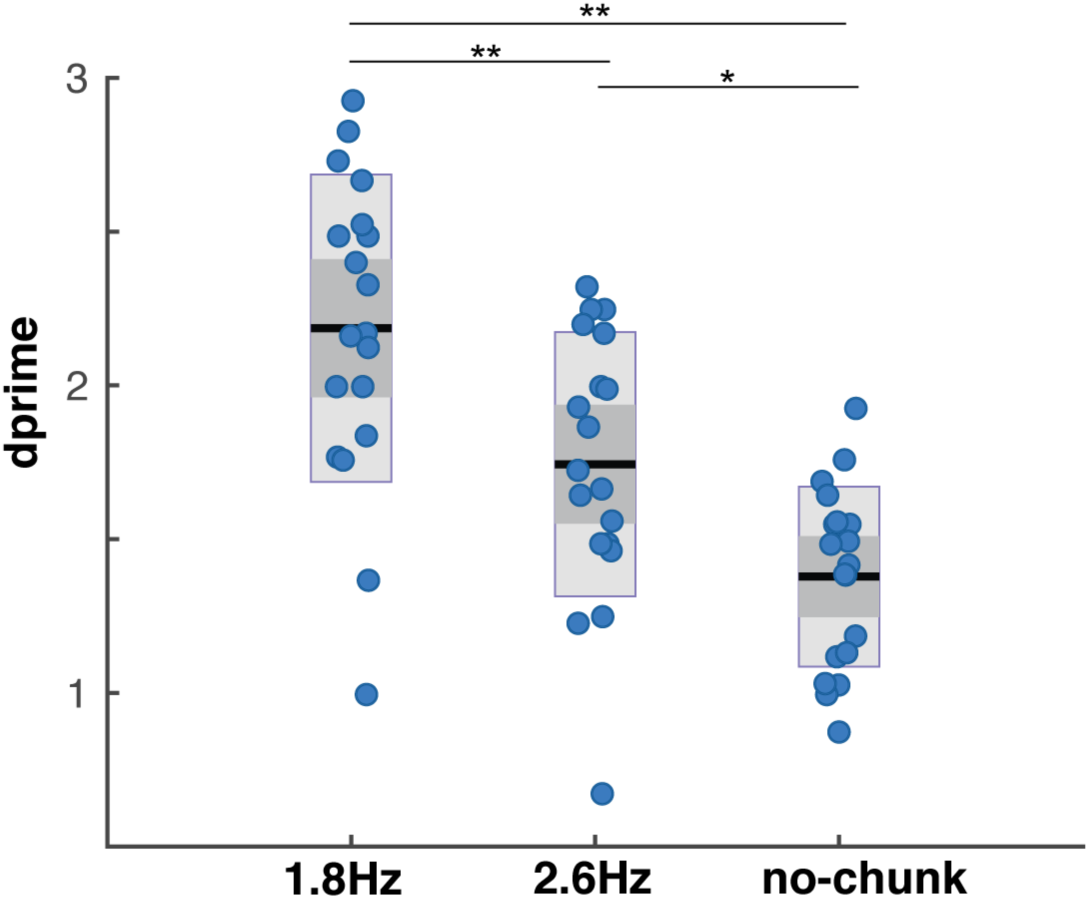
Behavioral performance in the digit retrieval task. Dprime values are displayed, as measure of performance accuracy, separately for each condition. Purple dots indicate individual dprime scores, red lines indicate the mean dprime scores, dark gray bars indicate the +/-1 standard error of the mean and light gray bars the confidence interval. Significance is indicated by *(p < .05) and **(p < .01). The performance was higher in the 1.8Hz acoustical chunk condition (inside delta chunking), compared to the 2.6Hz acoustical chunk (outside of delta) and the no-chunk condition (replicating findings by (Ghitza, 2017). The performance in the 2.6Hz was higher compared to the no-chunk condition.

### Probability density function (PDF) of periodicities of elicited brain waves

Fig. 2 shows the periodicity PDFs in the [1 4] Hz frequency range for the STG region of interest (ROI) in the left hemisphere. For each brain signal periodicity was extracted by a newly developed analysis method termed aggregated cross-correlation (XCOV), described in the Materials and Methods section. In constructing a histogram of periodicities, one XCOV peak in the [1 4] Hz frequency range was considered. A 3^rd^order Gaussian mixture model (GMM) that fits the histogram is the desired periodicity PDF. In Fig. 2, each entry shows the periodicity PDF; the “goodness” of the periodicity is quantified in terms of P – the percentage of datapoints inside the [1 4] Hz frequency range with respect to the total number of datapoints; and the mean μ and variance σ of the prominent Gaussian component of the 3^rd^ order GMM. The total number of data points is shown in the legend of each entry.

**Fig. 2.**
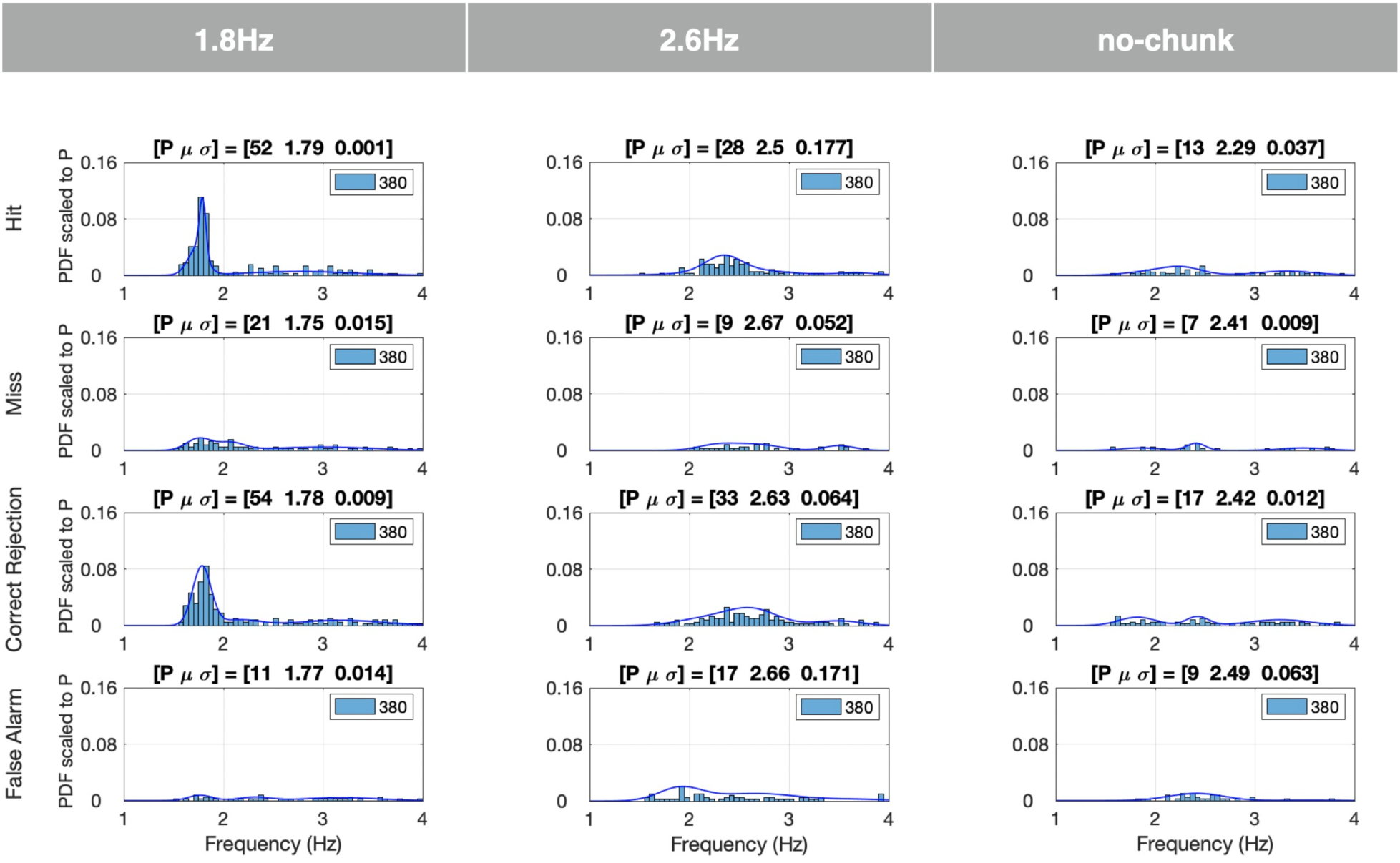
Probability density functions (PDFs) of delta periodicities per response class. The PDFs are displayed for the left hemisphere STG region of interest (ROI). The number of voxels in this ROI is 20 and the number of participants 19. Per voxel and subject, one XCOV peak inside the delta [1 4] Hz range was selected. The rows indicate the response classes (Hit, Miss, etc.), and the columns – the chunking conditions. Each entry shows the periodicity PDF, and the “goodness” of the periodicity is quantified in terms of a P value (defined in caption of Fig. 9), and the mean μ and variance σ of the prominent Gaussian component of a 3^rd^ order GMM. The Legend of each entry shows the total number of data points (20 voxels × 19 subjects). For the 1.8Hz condition, a strong periodicity presence at 1.8 Hz was recorded for the Hit and Correct Rejection responses, with P over 50%.. A much weaker presence was recorded for the Miss and False Alarm responses. A similar trend is shown for the 2.6Hz condition, albeit with much weaker periodicity presence compared to the 1.8Hz condition, and a smaller number of datapoints (P of about 30%).. Notably, no periodicity presence at 2.6 Hz was recorded for the no-chunk condition.

For the 1.8Hz condition, a strong periodic response at about 1.8 Hz was recorded for the Hits and Correct Rejections, with P over 50%. Much weaker presence of periodicity was recorded for the Misses and False Alarms. A similar trend is shown for the 2.6Hz condition, albeit with much weaker periodicity compared to the 1.8Hz condition, and with a smaller P (of about 30%). No periodicity at 2.6 Hz was evident for the no-chunk condition. Notice that, across chunk conditions, the PDF patterns for hits and correct rejections are similar, as are the patterns for misses and false alarms. Such similarities were observed for all ROIs. Therefore, in presenting the rest of the data, the hits and correct rejections are combined to indicate *Correct responses*, and the misses and false alarms are as *Erroneous responses*.

In the following figures, the data are presented as follows. Each figure contains 6×3 entries organized in six rows (ROIs) and three columns (chunking conditions). Each entry shows the periodicity PDF, and the “goodness” of the periodicity is quantified in terms of P, μ and σ Defined in Fig. 2. The legend of each entry is as defined in Fig. 2. In some selected entries, the upper left corner shows the Kullback-Leibler Divergence (KLD) of the entry’s PDF with respect to a reference PDF defined in the respective figure caption. Finally, in some entries, no μ and σ values are present. This is so because of a failure of the 3rd order GMM to converge due to the small P value.

Figures 3a and 3b show the elicited responses in the [1 4] Hz frequency band for Correct responses (i.e., Hits and Correct Rejections combined), and Erroneous responses (i.e., Misses and False Alarms combined), respectively. We term these elicited responses *delta responses*. For Correct responses in the 1.8Hz condition a strong periodicity presence at about 1.8 Hz is recorded. A similar pattern is shown for the 2.6Hz condition, albeit with much weaker periodicity presence compared to the 1.8Hz condition (lower P value and wider σ). No periodicity presence at 2.6 Hz is recorded for the no-chunk condition. For Erroneous responses, for all ROIs, no presence of periodicities is recorded, for any condition. More specifically: for Correct responses, in the chunked conditions, the auditory association ROI (STG) shows a compelling periodicity presence at 1.8 Hz in the 1.8Hz condition and a weaker presence at 2.6 Hz in the 2.6Hz condition. At the word-level ROI (MTG), periodicity exists for the chunked conditions, albeit with 1.8 Hz periodicity stronger than that of 2.6 Hz. In the speech-motor planning and integration ROIs (IFG, SMG, PC), periodicity is present at 1.8 Hz, albeit reduced, and is absent in the 2.6Hz condition. Note that in the visual ROI (Calcarine), delta periodicities are absent for all conditions. Importantly, a 2.6 Hz periodicities were absent in the no-chunk condition for Correct (and Erroneous) responses, for all ROIs. Finally, the 1.8Hz condition column of Fig. 3a also shows the Kullback-Leibler Divergence (KLD) for all ROIs, with respect to the STG ROI (highlighted in red). The KLD values suggest two distinct patterns of elicited delta periodicities, one observed in the ventral stream regions (STG and MTG ROIs, with KLD value of 0.032 for MTG), and the other in the dorsal stream regions (IFG, SMG and PC ROIs, with KLD values of 0.100, 0.162 and 0.316, respectively).

**Fig. 3.**
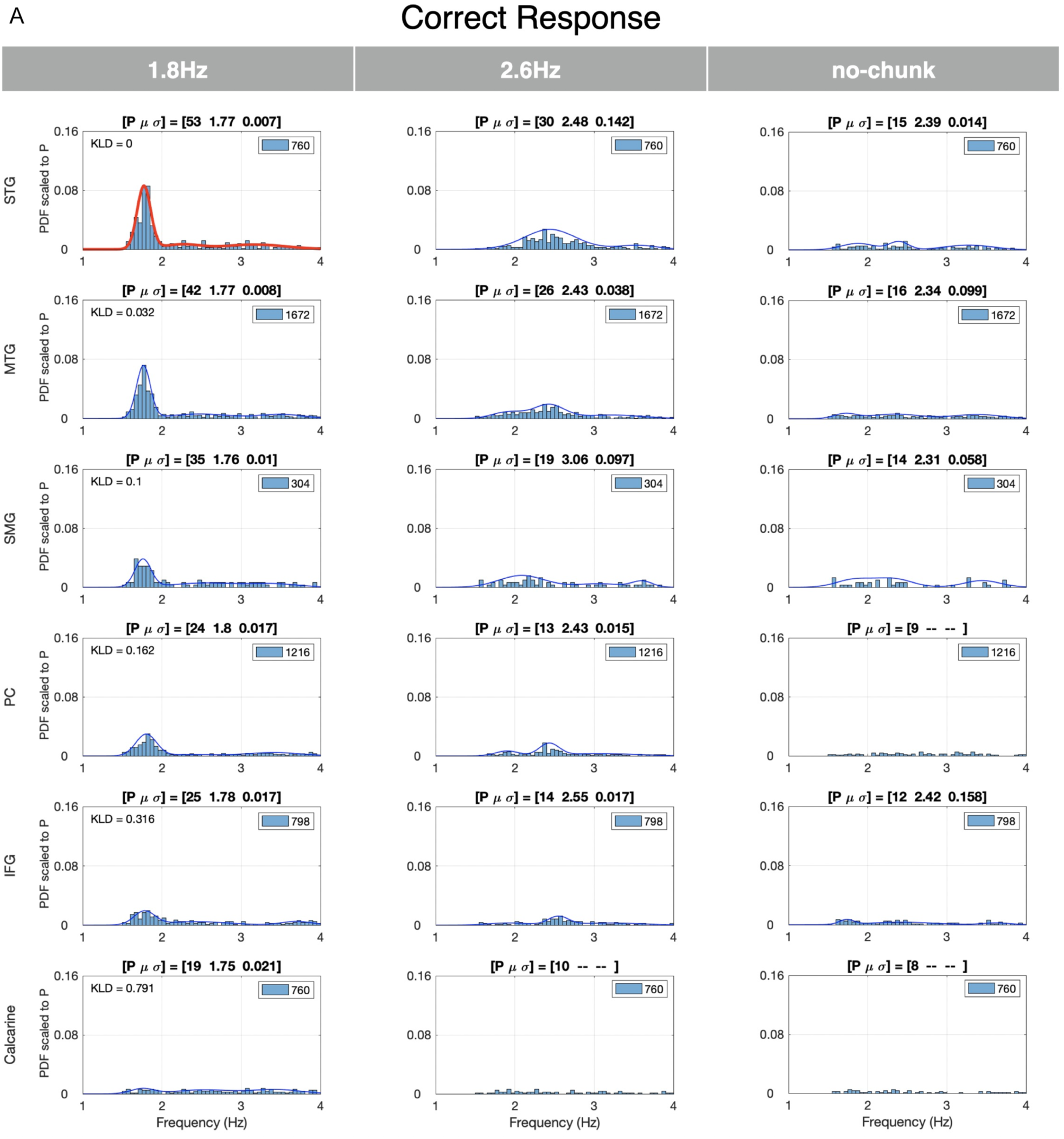
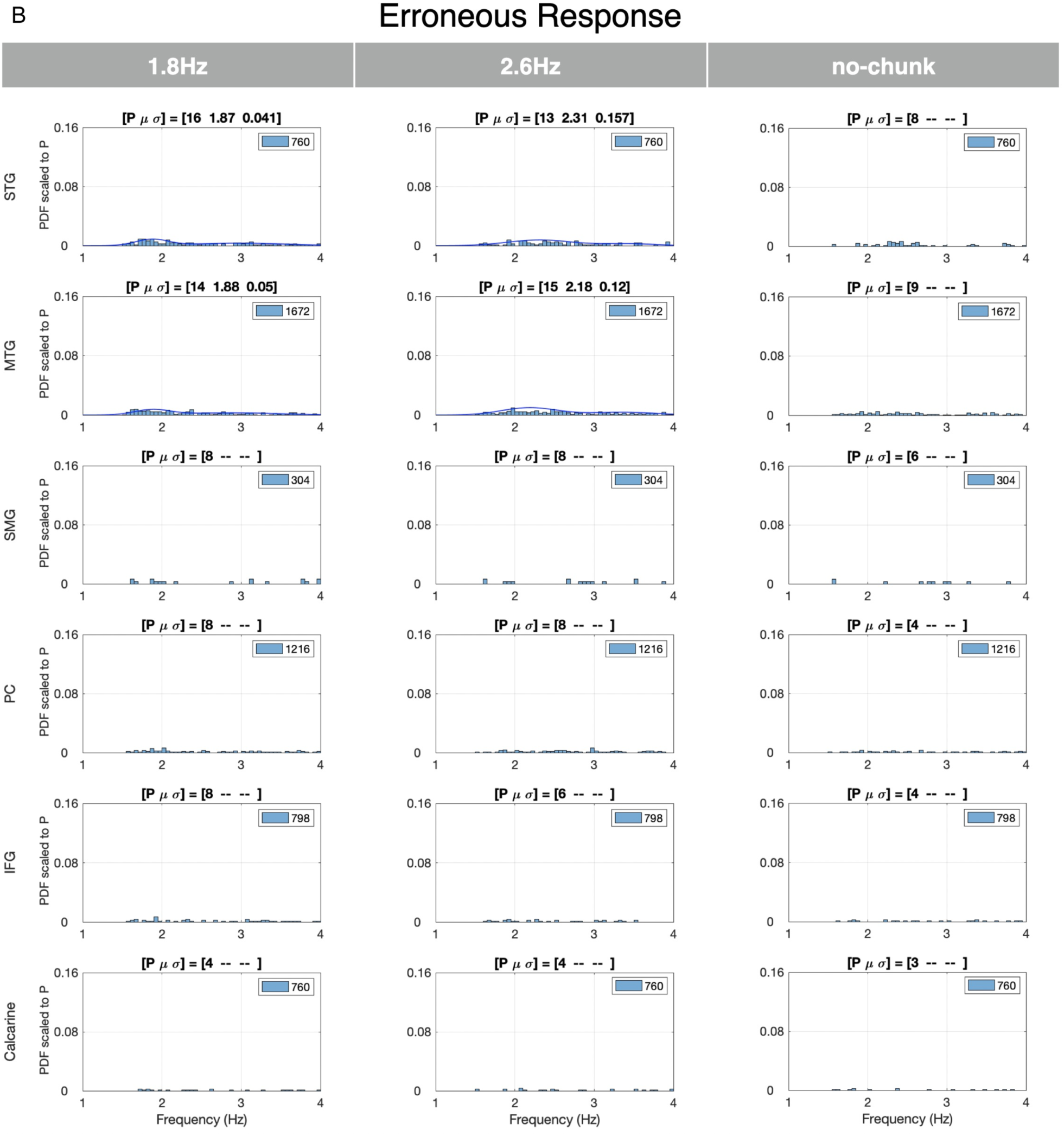
Delta periodicities for Correct and Erroneous responses in the left hemisphere. (a) Periodicities for Correct responses: Rows indicate the regions of interest (ROIs) and columns the chunking conditions. Each entry shows the periodicity PDF, quantified as defined in Fig. 4. For the 1.8Hz condition a strong periodicity presence at about 1.8 Hz is recorded. A similar trend is shown for the 2.6Hz condition, albeit with much weaker periodicity presence compared to the 1.8Hz condition. No periodicity presence at 2.6 Hz is recorded for the no-chunk condition. The 1.8Hz condition column shows the Kullback-Leibler Divergence (KLD) computed for this condition at all ROIs, with respect to the STG ROI highlighted in red (upper left corner of the ROIs). The KLD values suggest two distinct patterns of elicited delta periodicities, one observed in the ventral stream areas (STG and MTG ROIs), and the other in the dorsal stream areas (IFG, SMG and PC ROIs). (b) Periodicities for Erroneous responses: No presence of periodicities is recorded for any condition.

Figures 4a and 4b compare the elicited delta responses in all ROIs in the Left versus the Right hemispheres for Correct responses. Similar periodicity PDFs are observed for all ROIs in all chunking conditions. The 1.8Hz condition column of Fig. 4b shows the KLD of each ROI in the Right hemisphere, calculated against the corresponding Left ROI (Fig. 4a, highlighted in red). The KLD values show a closer similarity between the periodicity PDFs of the ventral stream regions (STG and MTG, with KLD values of 0.00 and 0.044, respectively), compared to that of dorsal stream regions (IFG, SMG, and PC, with KLD values of 0.224, 0.140 and 0.106, respectively).

**Fig. 4.**
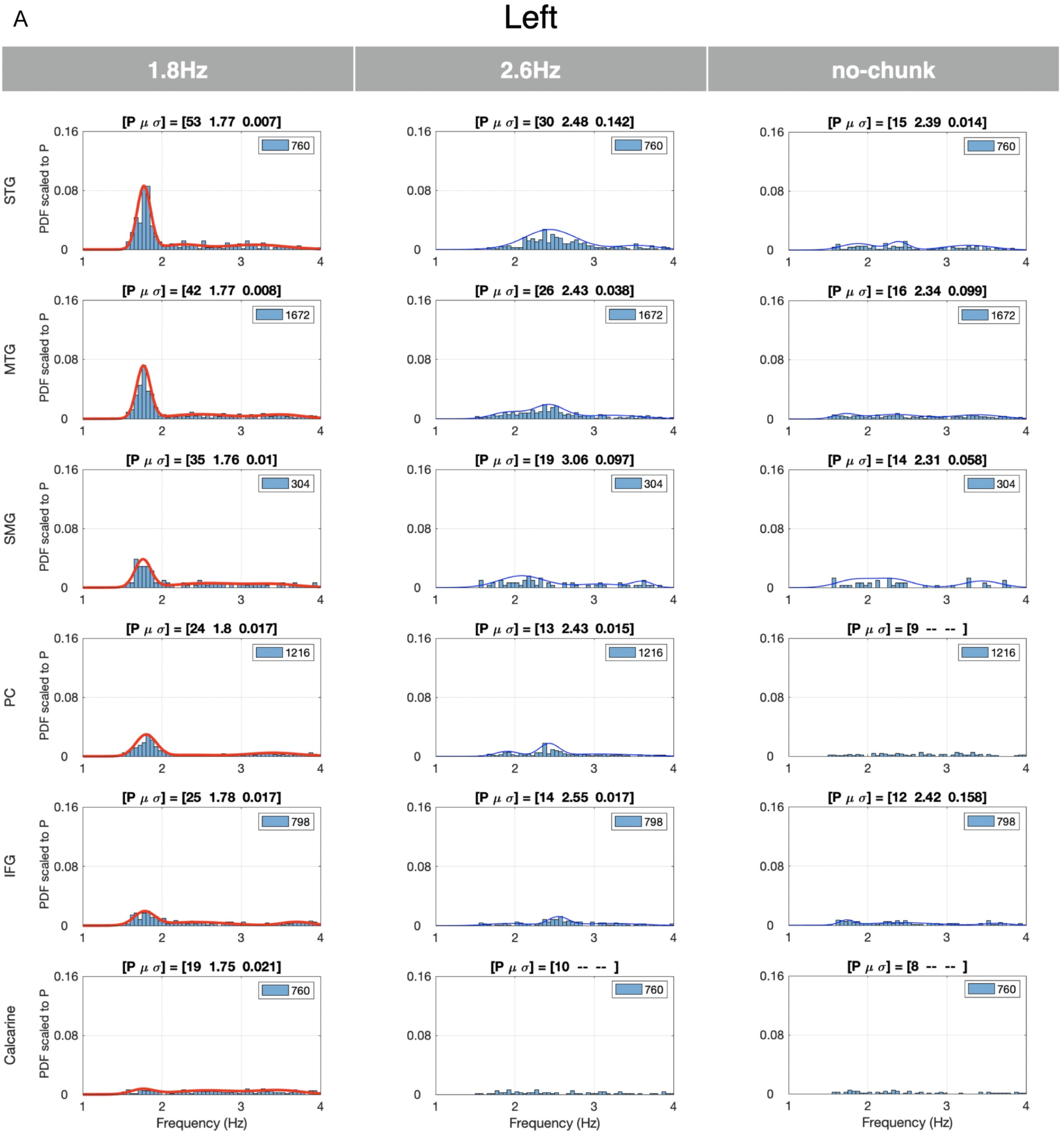
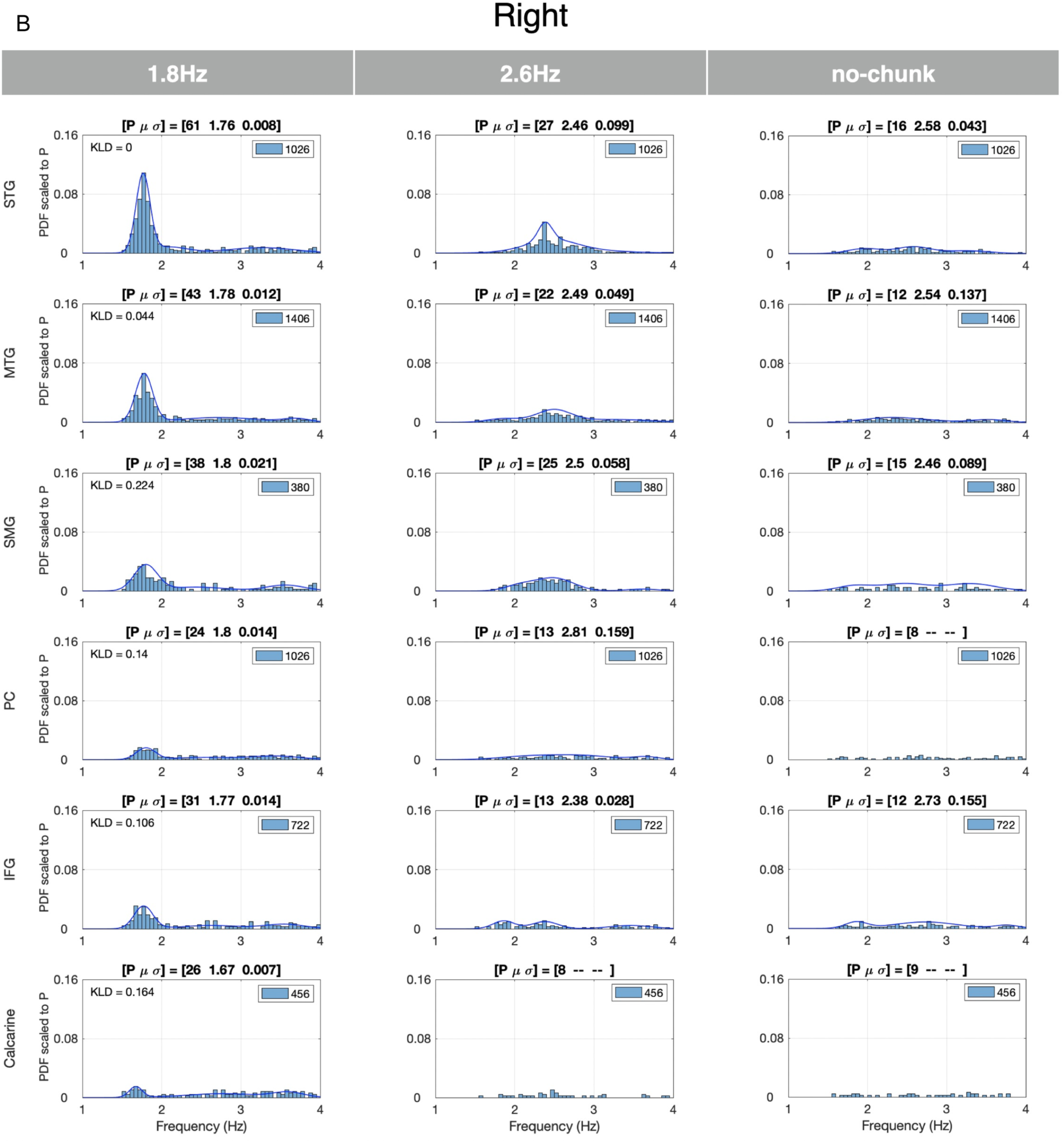
Comparing delta periodicities between the left and right hemispheres. (a) Shown are periodicity PDFs for the left hemisphere and Correct responses. They are compared with the periodicity PDFs for the Right hemisphere, shown in Fig. 7b. Similar PDF patterns are observed for all ROIs, in all chunking conditions. The 1.8Hz condition column of Fig. 7b shows the KLD of each ROI in the Right hemisphere, calculated against the corresponding Left ROI (Fig. 7a, highlighted in red). The KLD values show a closer similarity between the periodicity PDFs of the ventral stream areas (STG and MTG), compared to that of dorsal stream areas (IFG, SMG and PC). (b) Shown are periodicity PDFs for the right hemisphere and Correct responses. See caption of Fig. 7a.

Fig. 5a and 5b show the elicited responses in the [2 6] Hz frequency band for the Correct and Erroneous responses, respectively, for ROIs in the left hemisphere. We term responses in this frequency band *theta responses*. For the Correct behavioral responses, strong theta was elicited in all ROIs and for all chunking conditions, notably for the no-chunk condition. Such elicited neural response patterns reflect the single digit presentation rate. Two observations are noteworthy, the bimodal characteristic of the PDFs for all chunking conditions, in particular for the 1.8Hz chunking condition, and the strong, unexpected, theta periodicity presence in the Calcarine ROI. For the Erroneous responses, a weaker more dispersed periodicity presence was observed. Finally, for the Correct responses, the periodicity PDFs were similar in shape across conditions, as was quantified by the KLD values comparing the periodicity PDFs in the 1.8 Hz condition with respect to the 2.6 Hz condition (KLD values between 0.12 to 0.23 across ROIs) and the no-chunk condition with respect to the 2.6 Hz condition (KLD values between 0.01 to 0.10 across ROIs). The similarity of the PDFs across chunking conditions confirms that the decoding time at the digit level was sufficient across conditions.

**Fig. 5.**
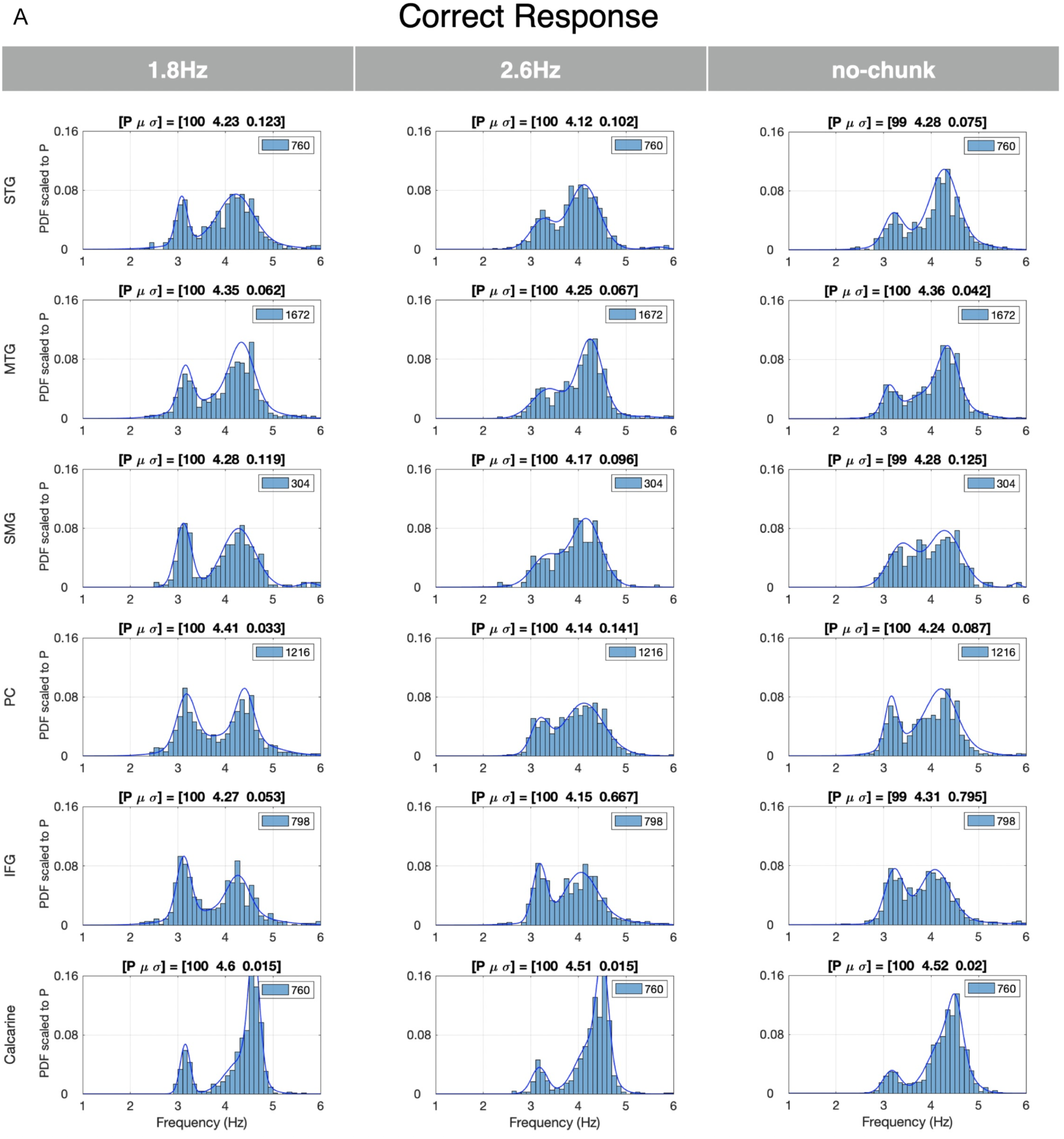
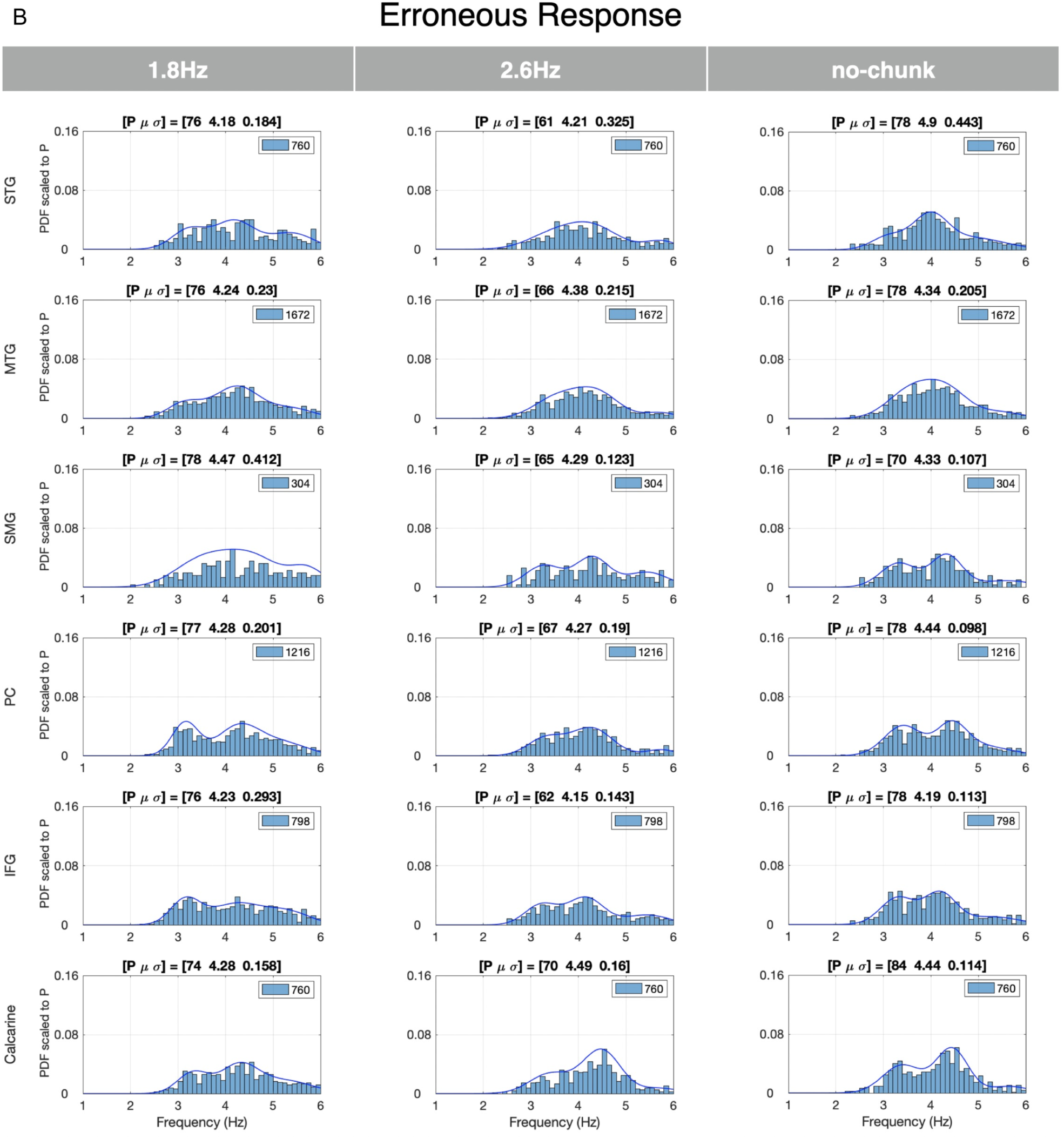
Theta periodicities for Correct and Erroneous responses in the Left hemisphere. (a) Periodicities for Correct responses: Strong theta periodicities were present in all ROIs and for all chunking conditions, notably also for the no-chunk condition. Such elicited neural response patterns reflect the single digit presentation rate. The PDFs exhibit a bimodal characteristic for all chunking conditions, in particular for the 1.8Hz chunking condition. (b) Periodicities for Erroneous responses: A weaker more dispersed presence of theta periodicities is recorded for all conditions (lower P value and wider σ).

### Correspondence between behavioral data and electrophysiological data

Fig. 6 quantifies the correspondence between the elicited delta periodicity patterns and the behavioral data (Fig. 1). Shown are the 3^rd^ order GMMs computed for the Correct responses in the left hemisphere and the three stimulus conditions (as in Fig. 3a). Unlike Fig. 3a, which shows PDF scaled to P, shown here are the actual PDFs (with the ∫*p(x)dx* = 1). The title of each panel shows three measures: (i) [*dprime* σ] – the behavioral performance indicated by mean *dprime* values and the variance across subjects (from Fig. 1); (ii) [*Bias* σ] – the average of the absolute difference (termed Bias) between the mean of the prominent Gaussian component of the GMM and the acoustic chunk rate, and the variance across the ROIs; and (iii) [*P* σ] – the average P value (defined in Fig. 2) and the variance across the ROIs. Two observations are noteworthy. First, the tightness of the PDFs in the 1.8Hz condition as reflected in the high probability value at the periodicity frequency, compared to the pseudo-uniform shape of the PDFs in the no-chunk condition. And second, the decrease in *dprime* is correlated with the increase in *Bias* and the decrease in *P*. These data support the hypothesis that perceptual chunking in the phrase level is derived by acoustic-driven delta oscillators.

**Fig. 6.**
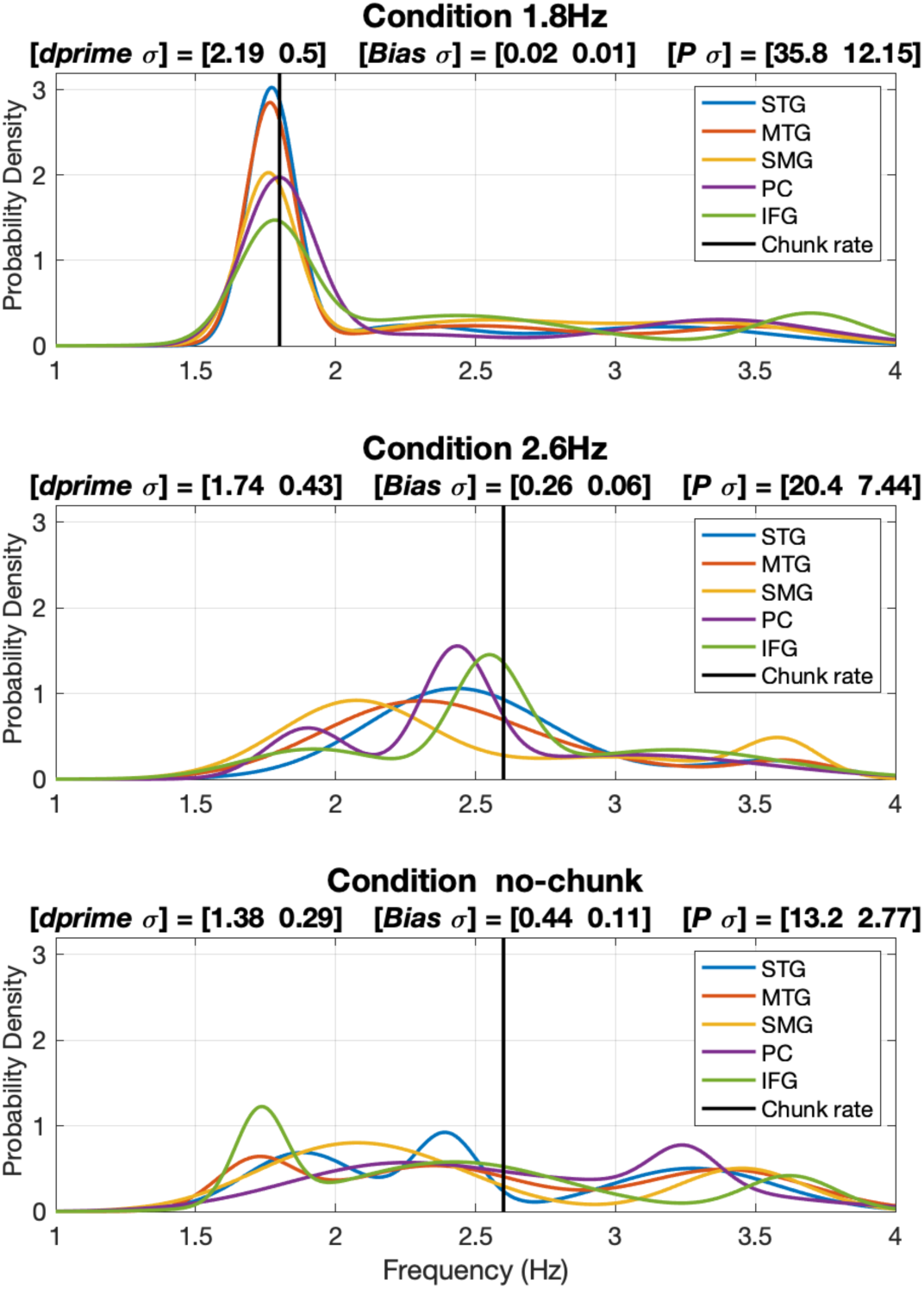
Correspondence between behavioral data and electrophysiological data. Shown are the 3rd order GMMs from Fig. 6a (i.e., Correct responses in the left hemisphere). Unlike Fig. 6a, which shows PDF scaled to P, shown here are the actual PDFs (with the ∫*p(x)dx* = 1). The title of each panel shows three measures: (i) [*dprime σ*] – the behavioral performance (from Fig. 2); (ii) [*Bias σ*] – the average of the absolute difference between the mean of the prominent Gaussian component of the GMM and the driving acoustic chunk rate, and variance across the ROIs; and (iii) [*P σ*] -- the average P-value (defined in Fig. 3) and variance across the ROIs. Note the tightness of the PDFs in the 1.8Hz condition compared to the pseudo-uniform shape of the PDFs in the no-chunk condition, and the correlation between the decrease in *dprime* and the increase in *Bias* and the decrease in *P*.

## Discussion

In this study we aimed to test, in electrophysiological terms, the hypothesis that the speech decoding process at the phrasal level is guided by a flexible, acoustic-driven neuronal delta oscillator locked to phrase-level acoustic cues (Ghitza, 2017). The proposal generalizes to the phrasal level the role played by neuronal theta-band oscillations in syllabic segmentation. We collected, concurrently, behavioral and MEG data during a digit retrieval task, in which digit sequences were either presented with an acoustic chunk pattern inside or outside of the delta range, or with no acoustic but a top-down invoked chunk pattern.

Stimuli with a chunk rate inside the delta range elicited considerable periodicity at the chunk rate in STG and MTG ROIs, and weaker periodicity at IFG, SMG and PC ROIs. Critically, this pattern of detected periodicities was directly related to Correct behavioral responses. In contrast, stimuli with a chunk rate outside of the delta range elicited weak periodicity, aligned with observed declines in behavioral performance. In the calcarine ROI (early visual cortex), considered a ‘control area’ for our analyses, no periodicities at the chunk rate were elicited. For the condition with no acoustic grouping but only the top-down induced chunk pattern (the no-chunk condition), periodicities at the demi-chunk rate were absent throughout the auditory pathway, for Correct and Erroneous responses.

Finally, a point that merits discussion. It could be argued that one cannot draw a conclusive relationship between ‘chunking’ and the neural signal in the delta range, because the 2.6Hz stimuli may be less intelligible due to the lack of silent gaps. Recall that the only difference between the 2.6Hz and the 1.8Hz stimuli is the insertion of a silent gaps in between the chunks (chunks being a digit doublet), but the chunks have identical time-compressed acoustics (see Fig. 7). At the digit level, strong theta periodicities (at the single digit rate) were elicited, with similar periodicity PDFs, at all ROIs, for all chunking conditions, and regardless of the level of behavioral performance (Fig. 5A). This data shows that the decoding time at the digit level was sufficient. In contrast, different periodicity patterns were observed at the chunk level, suggesting that the drop in behavioral performance for the 2.6Hz condition – with a chunk rate just outside the delta frequency range – is due to lack of chunk decoding time but not digit decoding time. Performance is recovered by bringing the chunk rate back inside the delta range by inserting gaps between chunks, hence providing the extra decoding time needed. As a whole, therefore, our data suggest that perceptual chunking at the phrasal level is realized by neuronal delta oscillators, and that the phrasal decoding time is determined by delta, in analogy to the role of theta in determining the decoding time at the syllable level (Ghitza, 2014).

**Fig. 7.**
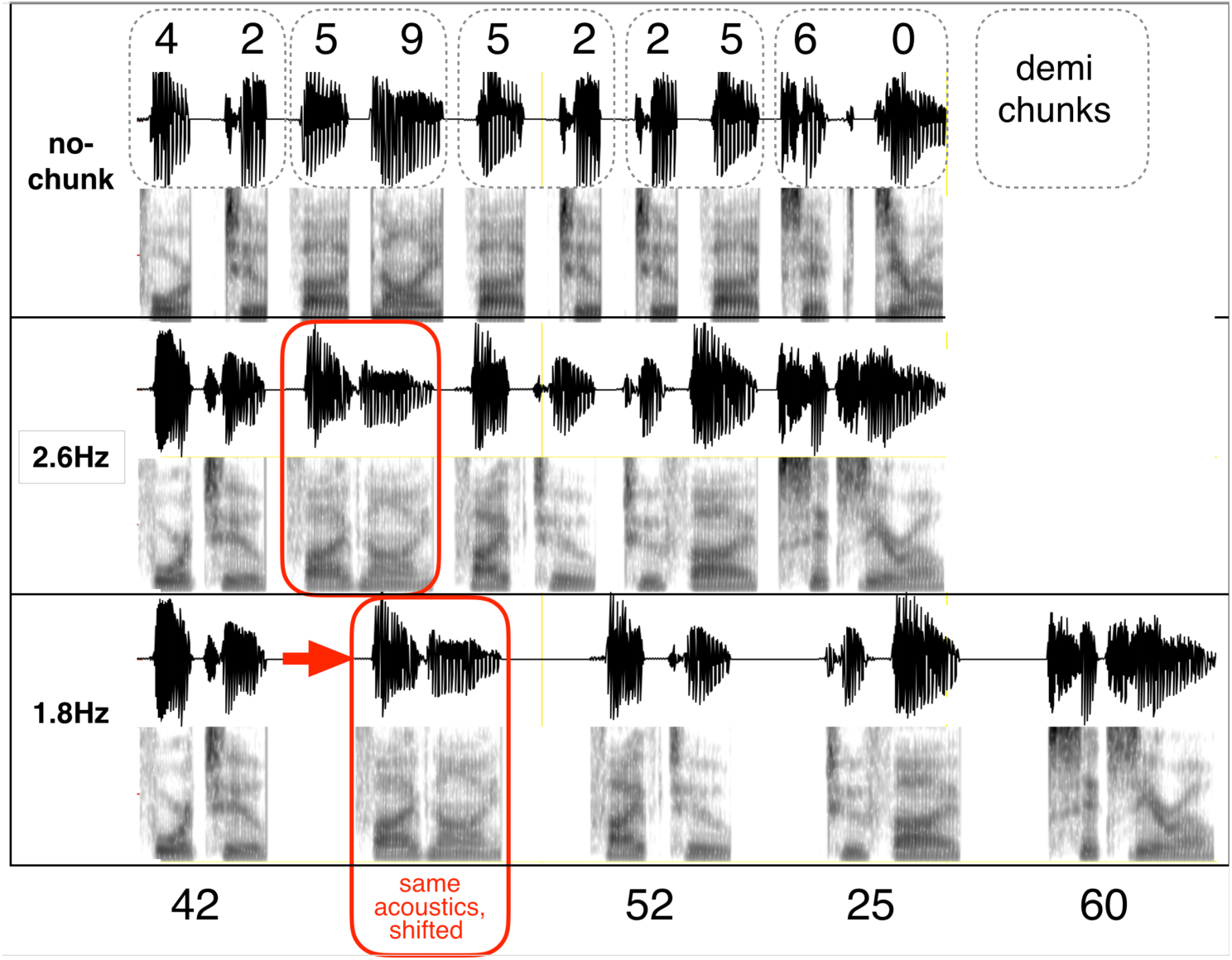
Chunk patterns and chunk rates for the 10-digit digit string 4259522560. (i) Lower two panels: These conditions were designed to address research-question-1. The chunk pattern is 2222, with chunk rates of 1.8 Hz (inside delta) and 2.6 Hz (outside). Each chunk was synthesized as a 2-digit unit, using the AT&T Text-to-Speech System accentuation (see text). Note that a particular 2-digit chunk has the same acoustics, regardless of whether it occurs in the 1.8Hz or 2.6Hz 2222 chunk condition (red box). The 1.8Hz stimulus is generated by increasing the gap between the chunks (with identical chunk acoustics). (ii) Upper panel: The “no-chunk” condition is illustrated, which was designed to address research-question-3. The single digit rate is 5.2 Hz. Although no acoustic cues for 2-digit chunking existed, a “cognitive 2222” pattern (with a demi-chunk rate of 2.6 Hz) is imposed on the acoustic 1111 stimuli by instructing the participants that the digit sequences they hear has a 2222 chunk pattern.

**Fig. 8.**
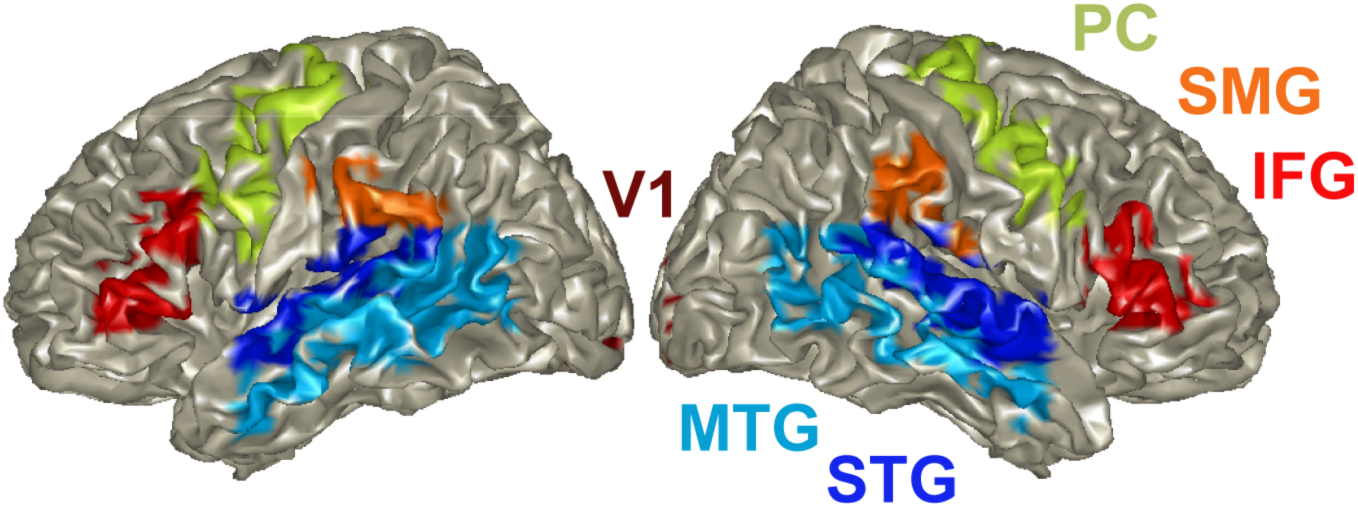
Cortical regions of interest (ROIs). The automated anatomical labeling atlas (AAL; Tzourio-Mazoyer et al., 2002) was used to select ROIs in left and right auditory cortex (STG), the ventral (MTG) and the dorsal (IFG, PC, SMG) auditory stream. V1 was used as control region. ROIs are color coded.

### Presence of delta periodicities in the ventral and dorsal stream

How should these activity patterns of neuronal delta and theta periodicities, be interpreted? In the ventral stream (STG and MTG), robust periodicities were recorded mainly by stimuli with a chunk rate inside the delta range, and only for Correct behavioral responses. Periodicities in these brain areas were present even for acoustic chunk rates at the edge of the delta range, albeit considerably weaker. Note that the observed lack of hemispheric lateralization in auditory cortex in our study is in line with previous reports on bilateral theta/delta activity elicited to more complex speech stimuli (Assaneo et al., 2019; Flinker et al., 2019). Interestingly, in contrast to the ventral stream, in the dorsal stream speech-motor integration areas more divergence between the left and right hemisphere was observed, whereas the left hemisphere more tightly followed the chunking rate compared to the right. While periodicities were overall weaker in the dorsal stream (IFG, SMG and PC), they were absent for acoustic chunk rates outside the delta range. Furthermore, compared to ventral stream brain areas, dorsal stream areas showed broader frequency distributions around the phrasal chunk rate. These findings suggest an important role for high-level auditory association areas including the ventral and dorsal stream in phrasal chunking. Importantly, and quite remarkably, the delta-band activity in these areas was fully aligned with behavioral performance (i.e. delta activity was only elicited in Correct, but not in Erroneous responses). Keitel et al. (2018) and Morillon et al. (2019) recently proposed that delta oscillations in the motor cortex are involved in temporal predictions, affecting speech processing in the auditory cortex at a phrasal scale. A possible interpretation of their findings through the lens of our results is that acoustic-driven segmentation at the phrasal level takes place in STG, and the recorded behavioral performance with respect to chunk rate is a consequence of the goodness of segmentation. When the chunk rate is inside the delta band, successful segmentation results in delta activities in SMG/PC/IFG speech-motor integration areas that may reflect decoding processes and possibly auditory-motor mapping related processes (Park et al., 2015). In contrast, chunk rates outside of the delta band result in bad segmentation in STG, and in turn suppressed periodicities in SMG/PC speech-motor integration areas due to unreliable decoding and audio-motor mapping. This interpretation is in line with another study (Donhauser and Baillet, 2020) that reports strong delta activity in STG when the speech input was ‘informative’, which may be the consequence of appropriate segmentation.

### Acoustic driven versus top-down driven delta periodicities

The intriguing question whether segmentation at the phrasal level is acoustic driven or context invoked is yet to be resolved. Here we address this question in a reductionist way, by limiting both the contextual information in our corpus (digit strings) and the complexity of the task (target ID task). To test whether segmentation is acoustic-driven or not we used the chunked stimuli, while the no-chunk stimuli were used to test context-invoked effects. Note that within the reduced scope of our study, the term ‘context-invoked’ is inappropriate because the degree of contextual information in digit strings is null. Instead, we use the term top-down-driven, to indicate the cognitive 2222 pattern of the no-chunk stimuli (with a demi-chunk rate of 2.6 Hz, imposed on the acoustic 1111 stimuli by explicitly instructing the participants that the digit sequences they hear have a 2222 chunk pattern; Fig. 7).

Our data show that the elicited delta-band activity is acoustically driven. For the chunked stimuli, and throughout the auditory cortical hierarchy, if a periodicity occurred its frequency corresponded to the acoustic chunk rate. The data also show no top-down-driven delta-band presence. For the no-chunk stimuli, throughout the auditory cortical hierarchy, an acoustic-driven activity of ∼ 4.5 Hz (the digit rate of the no-chunk signal) is present, but a top-down-driven activity of 2.6 Hz (the cognitive chunk rate) is not observed.

At the syllabic level, it has been shown that auditory cortex neuronal oscillations in the theta range are elicited without higher-level speech processing (Howard and Poeppel, 2010; Rimmele et al., 2015), and oscillations occur in sync with acoustic cues at the syllabic scale (Doelling et al., 2014; Oganian and Chang, 2019). At the phrasal level, knowledge about the nature of the acoustic cues that constitute accentuation is sparse, even though linguistic cues in the delta range have been pointed out to be relevant for comprehension (Aubanel et al., 2016). Two recent studies that examined the possible role of contextual information in eliciting cortical activity in the delta band merit discussion. Meyer et al. (2016) provided evidence for neuronal tracking of phrases based on linguistic cues, and suggested that syntax may override prosody-driven neuronal tracking of phrases. Ding et al. (2016) showed the existence of delta brain waves without any acoustic cues present at this linguistic level, suggesting that delta oscillations reflect the buildup of an internal linguistic structure. It is worth noting that the interpretation of Ding et al. is under debate, and that an alternative interpretation – not oscillation-based – have been suggested according to which the neuronal delta activity is driven by the isochronous nature of the driving stimuli. In summary, we conclude that the data patterns we observe reflect acoustic-driven neural activity in the relevant frequency band.

### Presence of theta periodicities in all chunking conditions

Our data show strong theta periodicities in all ROIs and for all chunking conditions, notably for the no-chunk condition. Such elicited neural response patterns reflect the single digit presentation rate. Two observations merit discussion. First, a bimodal characteristic of the PDFs is observed for all chunking conditions, but in particular for the 1.8Hz condition. The bimodality arises from the acoustic properties of the stimuli. Consider, for example, the stimulus shown in Fig. 7. For the no-chunk condition, the duration in between a digit and its neighboring digit is the same for all digits, however, since the digit duration varies among digits (‘one’ is shorter than ‘seven‘) the digit presentation rate exhibits a mild bimodal distribution. For the chunked conditions, three intra-digit durations can be identified: (i) the duration between the onset of the first digit of a chunk and the first digit in the following chunk, which gives rise to the chunking rate, (ii) the duration between the onset of the first digit and onset of the second digit in a chunk, and (iii) the duration between the onset of the second digit in a chunk and the onset of the first digit in the following chunk. This plurality in intra-digit durations give rise to a bimodal duration distribution with a skewness determined by the prescribed chunking rate. The skewness is accentuated, in particular, in our 1.8Hz stimuli. The bimodal nature in the acoustics drives the elicited neural response seen in our data (Fig. 5a).

The second observation of interest is the presence of strong theta periodicity, despite the absence of delta periodicity, in the calcarine ROI. Previous studies have shown that in audio-visual tasks, visual inputs related to visual stimuli have the potential to reset neuronal oscillations in auditory cortex if they are attended (Lakatos et al., 2009). Our data show action in the other direction, with acoustic driven theta oscillations present in visual areas (Luo et al., 2010). While several subcortical pathways for such supramodal effects were suggested, such direction of information flow could also be made possible through the motor cortex – a major nexus between auditory and visual paths. Indeed, in our data the presence of theta in the calcarine ROI, on the one hand, and the absence of delta there, on the other hand, is in correspondence with the elicited theta and delta activity patterns in the motor-cortex related ROIs (IFG, SMG and PC ROIs in Figs. 3a and 5a). Our explanation is in line with the fact that since eye movements can align to the timing of auditory stimulation (Joiner et al., 2007), and they do reset visual cortical activity (Barczak et al., 2019), phase reset related to saccadic eye movements link sensory cortical oscillatory activity across visual, motor and auditory regions.

### Oscillations versus evoked responses

Our data show strong delta cortical periodicities while listening to the 1.8 Hz chunked stimuli. Are these brain waves generated by a neuronal oscillator locked to the acoustic chunk rhythm or do they reflect the evoked response to the corresponding acoustic cues? The answer to this question at the syllabic level has been difficult to determine, because the impulse response of the neuronal circuitry to discrete acoustic cues associated with syllables (e.g., acoustic edges, vocalic nuclei) corresponds, in duration, to the theta-cycle range (about [125 330] msec). Doelling et al. (2019) addressed this conundrum by generating simulated outputs of an oscillator model and of an evoked response model, and comparing the quantitative predictions of phase lag patterns generated by the two models against recorded MEG data. They showed that, compared to the evoked response model, the model of oscillatory dynamics better predicted the MEG data. Our data provides additional support for the oscillator interpretation. Can the observed, robust periodic responses to a 1.8 Hz chunked stimulus reflect evoked responses elicited by discrete acoustic cues at the phrase level? Such a view requires a neuronal circuitry with an impulse response that is over half a second long in duration – a requirement at odds with known biophysical properties. Moreover, only a model of delta oscillatory dynamics can explain the fact that neural response at the delta range is present when the acoustic chunk rate is inside, but is absent for rates outside the delta range.

### Generalizability of the neuronal chunking mechanism

#### Scaling up to real speech

The studies discussed above (Meyer et al., 2016; Keitel et al., 2018; Morillon et al., 2019) suggest a presence of delta brain waves in phrasal chunking for continuous speech, beyond the digit retrieval paradigm used here. Extending our results to naturalistic speech has important implications for what would constitute optimally sized acoustic chunks for the sentential decoding – or parsing – process. If the information ‘bound’ within windows of roughly a delta cycle are integrated as phrases (intonation phrases and perhaps structural phrases, depending on the specific relation), it suggests that there are natural patterns of spoken phrase rhythms or phrase durations that are best suited for decoding spoken language, driven by the necessity to match a cortical function. Deploying the experimental analysis approach we describe here to real speech can elucidate the temporal aspects of spoken language comprehension.

#### Infra-delta chunking rate

The main research question of our study was whether elicited delta cortical oscillations correlate with behavior. In particular, does performance deteriorate if the chunk rate is outside the delta range? We addressed this question by looking at an above-delta chunking rate (2.6 Hz), but we didn’t look at infra-delta rates (e.g., 0.3 Hz). The reason to skip the effects of infra-delta rates stemmed from the fact that the decay time of echoic memory – about 2 sec long (e.g. (Cowan, 1984)) – roughly coincides with the lower bound of the delta-cycle duration. Consequently, the dominant factor at the origin of a possible deterioration in performance may very well be an internal time constraint on processing spoken material (due to echoic memory span) rather than prosodic segmentation.

## Conclusion

Oscillation-based models of speech perception postulate a cortical computational principle by which decoding is performed within a time-varying window structure, synchronized with the input on multiple time scales. The windows are generated by a segmentation process, implemented by a cascade of oscillators. At the pre-lexical level, the segmentation process is realized by a *flexible theta oscillator* locked to the input syllabic rhythm, where the theta cycles constitute the syllabic windows. Doelling et al. (Doelling et al., 2014) provided MEG evidence for the role of theta, showing that intelligibility is correlated with the existence of acoustic-driven theta neuronal oscillations.

Our major finding – that phrasal chunking is correlated with acoustic-driven delta oscillations – generalizes the role played by neuronal theta-band oscillations in syllabic segmentation to the phrasal level. At the phrase level, we suggest that the segmentation process is realized by a *flexible delta oscillator* locked to the input phrase-chunk rhythm, where the delta cycles constitute the phrase-chunk windows.

We argue that these findings can be generalized to continuous speech (i.e., beyond digit strings) and that intonational phrase patterns of language could be constrained by cortical delta oscillations. Adopting the view that the strategy of composing syllables and words into phrasal units is the result of an evolutionary trajectory to match a cortical function, we hypothesize that the phrases of language are constrained by delta oscillations, and the rules of chunking in speech production may be the product of common cortical mechanisms on both motor and sensory sides, with delta at the core.

## Materials and Methods

### Participants

The data from 19 healthy right-handed (Oldfield, 1971; mean score: 75.22, SD: 18.08) participants were included in the study (mean age: 24.89 years, SD: 3.54; f = 14). Two participants were excluded because of technical issues, and one participant because of outlier performance (i.e. performance < mean performance -2SD). Individual MRI scans were recorded for all except for two participants who did not fulfill the MRI scanning criteria. All participants gave written informed consent for participating in the study and received monetary compensation. The study was approved by the local ethics committee of the University Hospital Frankfurt (SG2-02-0128).

### Digit string stimuli

We used 10-digit long stimuli with a carefully crafted temporal structure. We opted for digit sequences – semantically unpredictable material (i.e., no context) – in order to minimize the bottom-up/top-down interaction that is in play in setting perceptual phrasal boundaries. The digit sequences were grouped into chunks, with two chunk patterns, termed 2222 or 1111 pattern. For example, the 2222 pattern of the sequence 4259522560 is [42 59 52 25 60], and the 1111 pattern is [4 2 5 9 5 2 2 5 6 0].

2222 pattern stimuli were generated as described in the ***Corpus*** subsection below, using two chunk rates: 1.8 Hz (inside the delta range) and 2.6 Hz (outside/marginal), termed conditions ‘1.8Hz’ and ‘2.6Hz’ (Fig. 7). A strong 1.8 Hz response to 1.8Hz stimuli, together with a diminishing 2.6 Hz response to 2.6Hz stimuli would indicate an acoustic driven delta response. 1111 pattern stimuli were used with a digit rate of 5.2 Hz (no-chunk controls). Importantly, a “cognitive 2222 syntax”, with a ‘demi-chunk’ rate of 2.6 Hz, was imposed on the acoustic 1111 pattern stimuli by instructing participants that the digit sequence they heard had a 2222 chunk pattern. This condition is termed a ‘no-chunk’ condition (Fig. 7). A 2.6 Hz response to no-chunk stimuli would indicate a top-down driven delta.

### Corpus

The text material comprised 100 10-digit long text strings. Stimuli were generated by using the AT & T Text-to-Speech System^1^ with the female speaker Crystal. To generate stimuli with a 2222 chunk pattern, we first created a 2-digit chunk vocabulary as follows. For each doublet of digits that exists in the 100 text strings, a naturally sounding 2-digit chunk waveform was generated (naturalness was obtained by the AT & T system accentuation rules) resulting in a chunk-vocabulary. For a given text string, a 2222 10-digit stimulus was generated by concatenating the corresponding five 2-digit chunk waveforms pulled from the chunk-vocabulary. The *chunk rate* was set by adjusting the gap duration in between two successive chunks, resulting in a stimulus with a temporal structure but without any contextual cues. For the 1111 chunk pattern, the same AT&T Text-to-Speech System was used to generate just 10 single-digit stimuli (0 to 9). The resulting vocabulary of single digits was used to generate any given 10-digit stimulus by the means of concatenation, with a 5.2 Hz chunk rate. The chunk rate was set by adjusting the gap duration in between two successive digits. To enable the generation of stimuli with chunk rates greater than the delta frequency upper bound (2.6 Hz), all waveforms were time compressed by a factor of 2.5, just below the auditory channel capacity (Ghitza, 2014). The duration of the 10-digit stimuli varied across conditions; for the 1.8Hz condition: mean = 2.61 sec (VAR = 85.6 msec); for the 2.6Hz condition: mean = 1.99 sec (VAR = 85.6 msec); and for the no-chunk condition: mean = 1.95 (VAR = 89.6 msec).

For each of the 300 10-digit stimuli (100 stimuli for each of the 1.8Hz, 2.5Hz, and no-chunk conditions) a trial was created by concatenating the following waveform sequence: [one digit trial-count] [20-msec long gap] [10-digit string] [500-msec long gap] [2-digit target], resulting in one concatenated waveform per trial with durations that varied across trials and conditions. The 300 trials were scrambled, and the resulting pool of trials was divided into blocks, 50 trials per block. A jittered intertrial interval of 3-4.5 sec was presented between trials. The behavioral response (yes/no) was entered in the intertrial interval (which follows the 2-digit target). Overall two different sets of stimuli were used.

### Task

Behavioral and MEG data were collected while participants performed a digit retrieval task, in the form of an adapted Sternberg target identification task (Sternberg, 1966) (target ID task from here on): listeners heard a 10-digit stimulus followed by a 2-digit long target, and were asked to indicate whether or not the target was part of the utterance. A target position was always within a chunk (acoustic or demi). Note that the task is suitable for probing the role of acoustic segmentation in a memory retrieval task: a successful yes/no decision depends on how faithful the recognized chunk objects are, generated by a decoding process that, by hypothesis, depends on the goodness of phrasal segmentation.

### Procedure and Paradigm

Participants were seated in front of a board for instructions in the MEG testing booth. Binaurally insert ear-plugs (E-A-RTONE Gold 3A Insert Earphones, Ulrich Keller Medizin-Technik, Weinheim, Germany) were used for stimulus presentation. Two button boxes (Current Designs, Inc.) were used to record participants’ responses. The Psychophysics Toolbox (Brainard, 1997) was used to run the experiment. During the experiment, on each trial participants fixated the screen center (fixation cross) while listening to the digit sequences. The sounds were presented at a comfortable loudness level (∼70 dB SPL), which remained unchanged throughout the experiment. Overall the experiment lasted about 2.5 hours, including preparation time, recording time, breaks, and post-recording questionnaires.

Participants were presented with the task requirements. Importantly, they were instructed that all sequences comprise concatenated chunks of two-digits (a “syntax” cue for the no-chunk condition). Prior to the experiment, all participants performed a short training of three trials (with feedback) in order to familiarize themselves with the stimuli and task. The MEG was recorded during the target ID task. Participants were asked to indicate by button press (yes/no response; with the response hand balanced across participants; yes-hand right: N = 12) whether or not the target was part of the preceded utterance.

## MRI and MEG Data Acquisition

A 3 Tesla scanner (Siemens Magnetom Trio, Siemens, Erlangen, Germany) was used to record individual T1-weighted MRIs. MEG recordings were performed on a 269-channel whole-head MEG system (Omega 2000, CTF Systems Inc.) in a magnetically shielded booth. Data were acquired with a sampling rate of 1200 Hz, online denoising (higher-order gradiometer balancing) and online low pass filtering (cut-off: 300 Hz) was applied. Continuous tracking of the head position relative to the MEG sensors allowed correction of head displacement during the breaks and prior to each file saving of a participant, using the fieldtrip toolbox (http://fieldtrip.fcdonders.nl) (Stolk et al., 2013).

## Analysis

### Behavioral Analysis

A “yes–no” model for independent observations was used to compute dprime (Green and Swets, 1966). Four classes of response are considered: (1) Hit: a “yes” response when the target chunk is present in the digit sequence, (2) Correct Rejection: a “no” response when the target chunk is absent, (3) Miss: a “no” response when the target chunk is present, and (4) False Alarm: a “yes” response when the target chunk is absent. Non parametric Wilcoxon signed-rank tests were used to test differences in the mean dprime across conditions. Bonferroni correction of multiple comparisons was applied.

### MRI Analysis

The FieldTrip toolbox (http://fieldtrip.fcdonders.nl) (Oostenveld et al., 2011) was used for the MRI and MEG data analyses. The standard Montreal Neurological Institute (MNI) template brain was used for participants where an individual MRI was missing. Probabilistic tissue maps (cerebrospinal fluid gray and white matter) were constructed from the individual MRIs. Next, a single shell volume conduction model (Nolte, 2003) was applied to retrieve the physical relation between sensors and sources. Between the individual T1 MRI and the MNI template T1 a linear warp transformation was computed. A 8 mm template grid, defined on the MNI template T1, was warped on the individual head space by inversely transforming it, based on the location of the coils during the MEG recording and the individual MRI. Next, based on the warped MNI grid and the probabilistic tissue map a forward model was computed, and applied for source reconstruction. This allowed aligning the grids of all participants to each other in MNI space for the across participants statistics.

### MEG Preprocessing

Line-noise was removed using bandstop filters (49.5-50.5, 99.5-100.5, two-pass; filter order 4) and the data were band-pass filtered off-line (0.1–100 Hz, Butterworth filter; filter order 4). A common semi-automatic artifact detection procedure was applied: for artifact rejection, the signal was filtered to identify muscular artifacts (band-pass: 110-140 Hz) or jump artifacts (median filter) and z-normalized per time point and sensor. The z-scores were averaged over sensors, in order to accumulate evidence for artifacts that occur across sensor. Trials that exceeded a predefined z-value (muscular artifacts, z = 15; jumps, z = 30) were rejected. Trials were the range (min-max difference) in any sensor exceeded a threshold (threshold = 0.75e-5) were identified as containing slow artifacts and rejected. Down-sampling to 500 Hz was applied. The data were epoched (−3.5 to 5 sec). Furthermore, when head movements exceeded a threshold (5 mm) a trial was rejected. Next, all blocks of recorded MEG data were concatenated. If high variance was detected at any sensor, the sensor was rejected. Finally, independent component analysis (infomax algorithm; Makeig et al., 1996) was used to remove eye-blink, eye-movement and heartbeat-related artifacts based on cumulative evidence from the component topography and time course.

### MEG source-level analysis

In a first step, the data were epoched (0-5 sec). For the main analyses only trials in which participants showed Correct responses (i.e. hits and correct rejections) were selected. Next, the sensor-space measurements were projected and localized in source-space inside the brain volume (Van Veen et al., 1997) using Linearly Constrained Minimum Variance (LCMV) beamforming. A spatial filter was computed based on the individual leadfields for each participant and condition (lamda = 10%; 0.8 cm grid). Next, all trials were epoched to the minimum stimulus duration in the corresponding condition (condition 1.8Hz: 2.38 sec; condition 2.6Hz: 1.68 sec; No-chunk condition: 1.75 sec).

### Cortical regions of interest (ROIs)

The automated anatomical labeling atlas (AAL; Tzourio-Mazoyer et al., 2002) was used to select the regions of interest (ROIs) as follows:

1. STG (Temporal_sup_L/R): Auditory association areas (Binder et al., 2009; Hickok and Poeppel, 2007)
2. MTG (Temporal_Mid_L) : Implicated in processing word form and meaning
3. IFG (Frontal_Inf_Tri_L/R): Involved in speech-motor planning
4. PC (Precentral_L/R), SMG (SupraMarginal_L/R): Speech-motor integration
5. Calcariane (Calcarine_L/R): Primary visual cortex (as a control region)

We opted to omit Heschl’s Gyrus (primary auditory cortex area) from the list of ROIs because of the very small number of voxels (3 in the Left, 2 in the Right).

### Probability density function (PDF) of periodicities within ROI

We aim to determine whether the elicited brain signal measured at any given voxel within a specific ROI shows periodicity, and if so, to extract the frequency. Ultimately, we seek to characterize the probability density function of the periodicities across all voxels in the ROIs of interest.

#### The aggregated cross-correlation measure (XCOV) of periodicity

To measure the neural response periodicity in individual voxels we used a newly developed analysis method reasoned as follows. Assuming that the number of trials is M one could calculate, e.g., the intertrial linear coherence (ITLC) or the intertrial phase coherence (ITPC). The outcome would be the frequency distribution of the coherence function, based on the average across the M trials. Another way to measure periodicity is by autocorrelation, where the first nontrivial peak indicates the period. Importantly, these measures build on the number of trials M. The trial signals are noisy, both due to the SNR and due to the brain wave irregularity – which is why these methods average over trials. What if M is too small? We propose a new method, termed ‘Aggregated cross-correlation’ (abbreviated XCOV), to measure periodicity across M trials. Broadly, we suggest to take advantage of the fact that, for M trials, we can generate about M^2^/2 cross-correlation functions. Note that, unlike autocorrelation, the first peak of a cross-correlation function does not indicate the period but rather the delay between the two signals. Therefore, we run each of the M^2^/2 candidate pairs through a “match filter”, which determines whether the corresponding two signals have a “zero” delay. Such a pair will have a cross-correlation function similar to that of an autocorrelation function, i.e., its peak is at zero and its earliest nontrivial peak is at the period. Only the pairs that pass the test are cross-correlated and aggregated. Obviously, the number of cross-correlation functions qualified for aggregation is between M and M^2^/2, depending on how strict the match filter is. We term the outcome of this procedure as the ‘XCOV’ function.

#### Periodicities PDF within a ROI

Fig. 9 details the analysis pipeline for deriving the probability density function (PDF) of the periodicities within a particular ROI. L voxels, N subjects, and M trials per subject are considered. First (not shown), each brain signal is filtered to the frequency range of interest (low pass filter with cutoff frequency of 6 Hz for delta analysis and a bandpass filter with a [2-10] Hz frequency range, for theta analysis). Cross correlations were computed using the filtered signals. Shown is the XCOV function at the i-th voxel, for the j-th subject, obtained by aggregating K cross-correlation functions. (Note that as a cross-correlation function, XCOV is computed against time-lags; the abscissa here shows the time-lag *inverse*, in frequency, hence going from right to left). The particular XCOV function shown has a single peak at 1.76 Hz but note that, in general, an XCOV may have more than one local peak. Next, the location of the prominent peaks^2^ are extracted, with the number of prominent peaks as a parameter. In our analysis one prominent peak per XCOV is considered. Hence, for L voxels and N subjects, a maximum of L×N data points are available to construct a histogram, from which only those inside the frequency range of interest are used, and the resulting histogram is normalized to L×N. A 3^rd^ order Gaussian mixture model (GMM) that fits the histogram is the desired PDF. The “goodness” of the periodicity is quantified by in terms of P, the percentage of datapoints inside the frequency range of interest with respect to the total number of datapoints (L×N); and the mean μ and variance σ of the prominent Gaussian component of a 3rd order GMM. (The total number of data points is shown in the legend of each entry.)

**Fig. 9.**
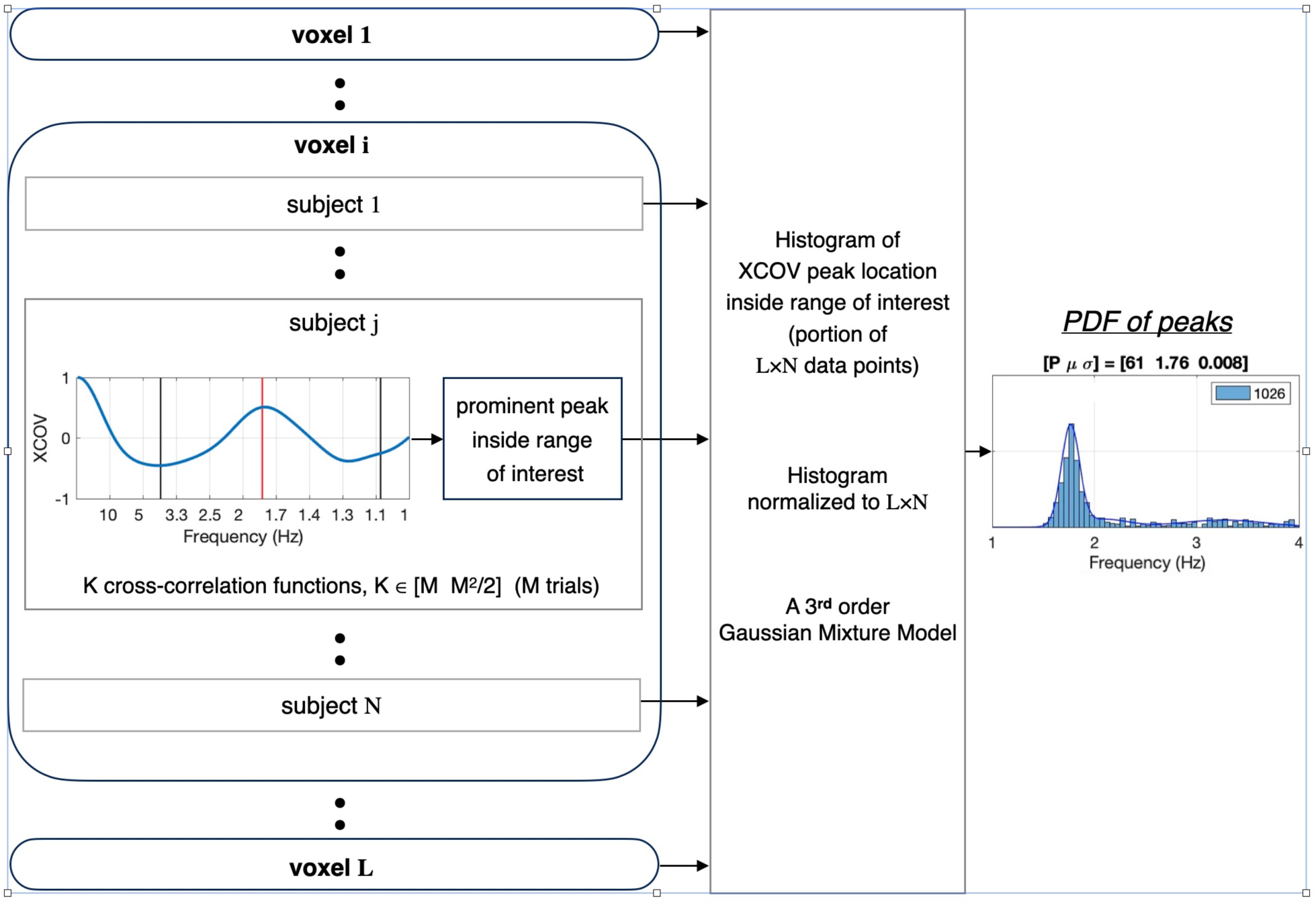
Analysis pipeline for deriving the probability density function (PDF) of the periodicities within a particular ROI. Shown is the resulting periodicity PDF for a given condition (say, the 1.8Hz condition), a given response class (say, Hit), and a given ROI (say, STG). L voxels, N subjects, and M trials per subject are considered. For the i-th voxel and the j-th subject, periodicity is computed using a newly developed analysis method termed ‘aggregated cross-correlation’, abbreviated XCOV. (Note that as a cross-correlation function, XCOV is computed against time-lags; the abscissa here shows the time-lag *inverse*, in frequency, hence going from right to left). A histogram of the XCOV periodicities is created and a 3^rd^ order Gaussian mixture model (GMM) that fits the histogram is the desired PDF. The “goodness” of the PDF is quantified by in terms of P value, the percentage of datapoints inside the frequency range of interest with respect to the total number of datapoints (L×N); and the mean μ and variance σ of the prominent Gaussian component of a 3^rd^ order GMM. (The total number of data points is shown in the legend of each entry.) (See text for more details).

## Acknowledgements

This work was funded by the Max-Planck-Institute for Empirical Aesthetics and by a research grant from the US Air Force Office of Scientific Research. We thank Simone Franz and Daniela van Hinsberg for help with the data recording.

## Footnotes

1 The AT&T-TTS system (http://www.wizzardsoftware.com/text-to-voice.php) uses a form of concatenative synthesis based on a unit-selection process, where the units are cut from a large, high-quality, pre-recorded natural voice fragments. The system produces natural-sounding, highly intelligible spoken material with a realistic prosodic rhythm––with accentuation defined by the system-internal prosodic rules––and is considered to have some of the finest quality synthesis of any commercial product.

2 The prominence of a peak measures how much the peak stands out due to its intrinsic height and its location relative to other peaks in the range of interest.

